# Kinesin-6 Klp9 plays motor-dependent and -independent roles in collaboration with Kinesin-5 Cut7 and the microtubule crosslinker Ase1 in fission yeast

**DOI:** 10.1101/476754

**Authors:** Masashi Yukawa, Masaki Okazaki, Yasuhiro Teratani, Ken’ya Furuta, Takashi Toda

**Affiliations:** Hiroshima Research Center for Healthy Aging (HiHA); Laboratory of Molecular and Chemical Cell Biology, Department of Molecular Biotechnology, Graduate School of Advanced Sciences of Matter, Hiroshima University, 1-3-1 Kagamiyama, Higashi-Hiroshima, Hiroshima 739-8530, Japan; Advanced ICT Research Institute, National Institute of Information and Communications Technology, Kobe, Hyogo 651-2492, Japan

## Abstract

Bipolar mitotic spindles play a critical part in accurate chromosome segregation. During late mitosis, spindle microtubules undergo drastic elongation towards the cell cortex in a process called anaphase B. Two kinesin motors, Kinesin-5 and Kinesin-6, are thought to generate outward forces to drive spindle elongation, and the microtubule crosslinker Ase1/PRC1 maintains structural integrity of antiparallel microtubules. However, how these three proteins orchestrate this process remains unknown. Here we explore the functional interplay among fission yeast Kinesin-5/Cut7, Kinesin-6/Klp9 and Ase1. Using total internal reflection fluorescence microscopy, we show that Klp9 is a processive plus end-directed motor. *klp9Δase1Δ* is synthetically lethal. Surprisingly, this lethality is not ascribable to the defective motor activity of Klp9; instead, it is dependent upon an NLS and coiled coil domains within the non-motor region. We isolated a *cut*7 mutant (*cut7-122*) that displays temperature sensitivity only in the absence of Klp9. Interestingly, *cut7-122* is impaired specifically in late mitotic stages. *cut7-122klp9Δ* double mutant cells exhibit additive defects in spindle elongation. Together, we propose that Klp9 plays dual roles during anaphase B; one is motor-dependent that collaborates with Cut7 in force generation, while the other is motor-independent and ensures structural integrity of spindle microtubules together with Ase1.

## INTRODUCTION

Bipolar spindle assembly is essential for proper sister chromatid segregation. Aberrations in this process result in chromosome missegregation and the emergence of aneuploid progenies, leading to miscarriages, birth defects and several human diseases, including cancer ^1, 2^. The formation of bipolar spindle microtubules consists of multiple, sequential steps, in which a myriad of proteins, including motor molecules, non-motor microtubule associated proteins (MAPs) and regulatory protein-modifying enzymes, act in a coordinated spatiotemporal manner. The establishment of spindle bipolarity is in sync with the physical separation of duplicated centrosomes (the spindle pole bodies, SPBs, in fungi), which is driven by the Kienisn-5 motor (budding yeast Cin8 and Kip1, fission yeast Cut7, *Aspergillus* BimC, *Drosophila* Klp61F, *Xenopus* Eg5 and human Kif11) ^3–9^. Accordingly, in many organisms, Kinesin-5s are essential for cell proliferation; their inactivation results in the formation of lethal monopolar spindles with unseparated centrosomes ^10–12^. Kinesin-5s form homotetramers, thereby crosslinking and sliding apart antiparallel microtubules ^13, 14^.

Interestingly, in many systems, outward forces generated by Kinesin-5s are antagonised by opposing inward forces elicited by Kinesin-14 motors; monopolar phenotypes resulting from inactivation of Kinesin-5s are rescued by simultaneous inactivation of Kinesin-14s ^15–24^. In fission yeast, temperature sensitive (ts) *cut7* mutations are suppressed by the deletion of either *pkl1* or *klp2* that encodes Kinesin-14 ^18, 19, 24-29^.

Once sister chromatids segregate towards each pole (anaphase A), during which spindle length is kept constant, spindle microtubules resume lengthening (anaphase B), followed by cytokinesis. Anaphase B proceeds through the sliding apart of antiparallel microtubules that generates outward force towards the two centrosomes ^30, 31^. Kinesin-5s are in general thought to be involved in anaphase B spindle elongation. However, as this kinesin is required for an initial step in the establishment of spindle bipolarity, a simple inactivation of Kinesin-5s leads to monopolar spindle formation without reaching anaphase. Therefore, it is not straightforward to prove the role of this kinesin in later stages of mitosis. In addition, there is another kinesin subfamily that plays a role in spindle elongation specifically during late mitotic stages: that is Kinesin-6 ^32, 33^.

Kinesin-6 family members (fission yeast Klp9, *C. elegans* ZEN-4, *Drosophila* Pavarotti and Subito, and human MKLP1/CHO1/Kif23, MKLP2/Kif20A/Rab6-KIFL and MPP1/Kif20B) are localised to the spindle midzone during anaphase B and form homotetramers like Kinesin-5s ^34–40^. In higher eukaryotes that possess multiple Kinesin-6 members (eg. human beings and fly), each member makes a distinct contribution to anaphase B spindle elongation and/or cytokinesis in a both independent and cooperative manner ^35, 41-44^. By contrast, in fission yeast, the sole member of Kinesin-6, Klp9, appears to play multiple roles on its own ^26, 37, 45^. Another crucial factor for anaphase B spindle elongation is the microtubule crosslinker PRC1/Ase1, which bundles antiparallel microtubules and ensures structural integrity of the spindle midzone ^46–50^. Whether Kiensin-5 or Kinesin-6 molecules play any role in microtubule bundling like PRC1/Ase1 independent of their motor activities remains unknown and therefore, is an important issue to be clarified.

We and others previously implicated that Klp9 executes essential functions for cell viability in a redundant fashion with Ase1 ^25, 26, 37, 45^; however, it is not understood as to how this kinesin functionally collaborates with Ase1. In addition, it has not been determined whether Cut7 acts in spindle elongation during late mitosis as well as in early mitosis. In this study, we address the following three specific questions. First, is Klp9 indeed a plus-end motor and if so, how does it behave in vitro? Second, how do Klp9 and Ase1 collaborate in spindle integrity during anaphase B? Lastly, does Cut7 drive spindle elongation during anaphase B as a motor? We show that Klp9 plays a microtubule-crosslinking role that is independent of plus end-directed motility and that Cut7 drives spindle elongation during anaphase B. Consequently, we can draw a detailed picture of the molecular interplays between two mitotic kinesins and the microtubule crosslinker during late mitotic stages.

## RESULTS

### Klp9 is a plus end-directed processive motor

Klp9 belongs to the kinesin-6 family, but its biophysical characterisation with regards to motor activity has not been reported. Therefore, as a first step, we expressed and purified recombinant full-length Klp9 protein tagged with EGFP, FLAG and 8xHis (Klp9-EGFP) (Supplementary Fig. S1) and performed in vitro assay with total internal reflection fluorescence microscopy (TIRFM). Purified Klp9 was immobilised onto a glass surface via an anti-His antibody (Fig. 1A), followed by the addition of paclitaxel (taxol)-stabilised, polarity-marked microtubules. This microtubule gliding assay showed that Klp9 could translocate microtubules with the minus end leading at a velocity of 39.0 ±0.3 μm/s (n=96, repeated 3 times) (Fig. 1B and 1C), indicating that Klp9 moved towards the microtubule plus ends.

**Figure 1.**
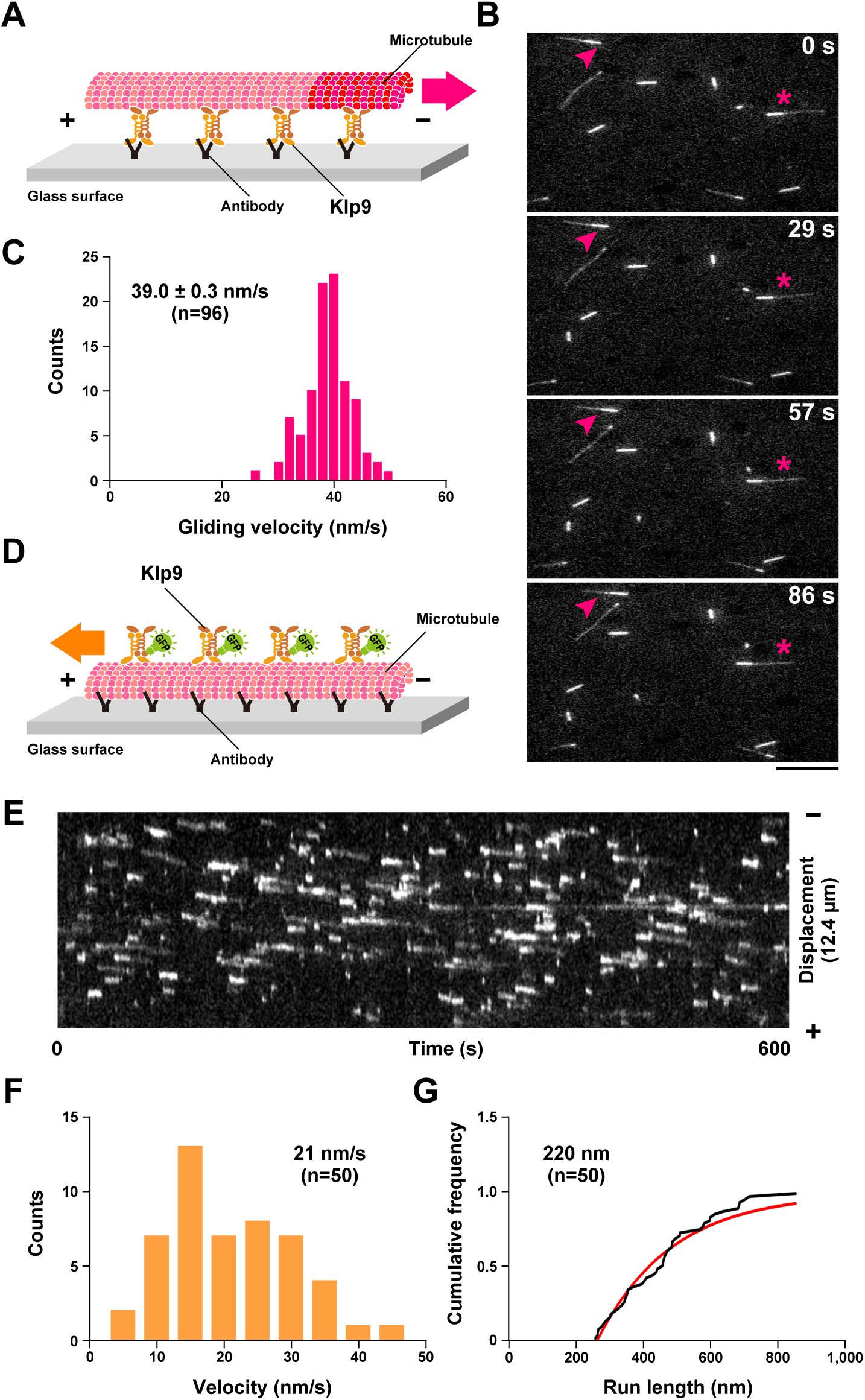
Klp9 is a processive plus end-directed motor. (A) Scheme of the in vitro microtubule gliding assay. (B) Representative time lapse images of the microtubule gliding assay. Paclitaxel-stabilised, polarity-marked microtubules were added to slides coated with purified Klp9 proteins in the presence of 1 mM ATP. Microtubule gliding was viewed by TIRFM. Images were taken every 1 s, and the time in the panels, in seconds, is shown on the top right. Two microtubule ends moving towards the minus ends are shown with arrowheads and asterisks. Scale bar, 10 µm. (C) Distribution and the average value of gliding velocity. (D) Scheme of the single-molecule assay. (E) Processive motility towards the plus end of paclitaxel-stabilised microtubules. Representative TIRFM kymograph depicting 0.3 nM Klp9-msfGFP in the presence of 1 mM ATP is shown. (F) Distribution and the average value of instantaneous velocity. (G) Cumulative run length. The distance between the appearance and disappearance of a fluorescent spot on the microtubule was measured.

Next, we added purified Klp9-EGFP proteins to immobilised, polarity-marked microtubules at a concentration of 0.3 nM and observed the behaviour of Klp9 at the single molecule level on the microtubule (Fig. 1D). As shown in Fig. 1E, Klp9 displayed processive motility toward the plus ends of microtubules. Quantification of Klp9 motility showed that Klp9 walked on the microtubule with a velocity of 21 nm/s (Fig. 1F, n=50) and its average run length was 220 nm (Fig. 1G, n=50). These in vitro results firmly establish that Klp9 is a processive motor with plus end-directed motility.

### The Klp9 motor is an important determinant for spindle elongation rate during anaphase B

We previously isolated a *klp9* ts mutant that contains a missense mutation within the N-terminal motor domain (*klp9-*2, K333M) ^26^. In addition, an ATPase-defective *klp9* rigor mutant containing a point mutation in the switch II region within the motor domain (Klp9^rigor^, G296A) was previously constructed ^45^. We examined Klp9 localisation (tagged with YFP under the native promotor) in these mutants and compared it with wild type Klp9 (a microtubule was simultaneously visualised with mCherry-Atb2 (α2-tubulin) ^51^). Interestingly, while Klp9-YFP concentrated on the middle region of spindle microtubules during anaphase B, corresponding to the spindle midzone in which antiparallel microtubules overlap ^37^, both Klp9-2-YFP and Klp9^rigor^-YFP were localised along spindle microtubules in a more diffused manner (Fig. 2A and 2B). These localisation patterns are consistent with the notion that Klp9 possesses a plus-end motility, thereby accumulating on the spindle midzone.

**Figure 2.**
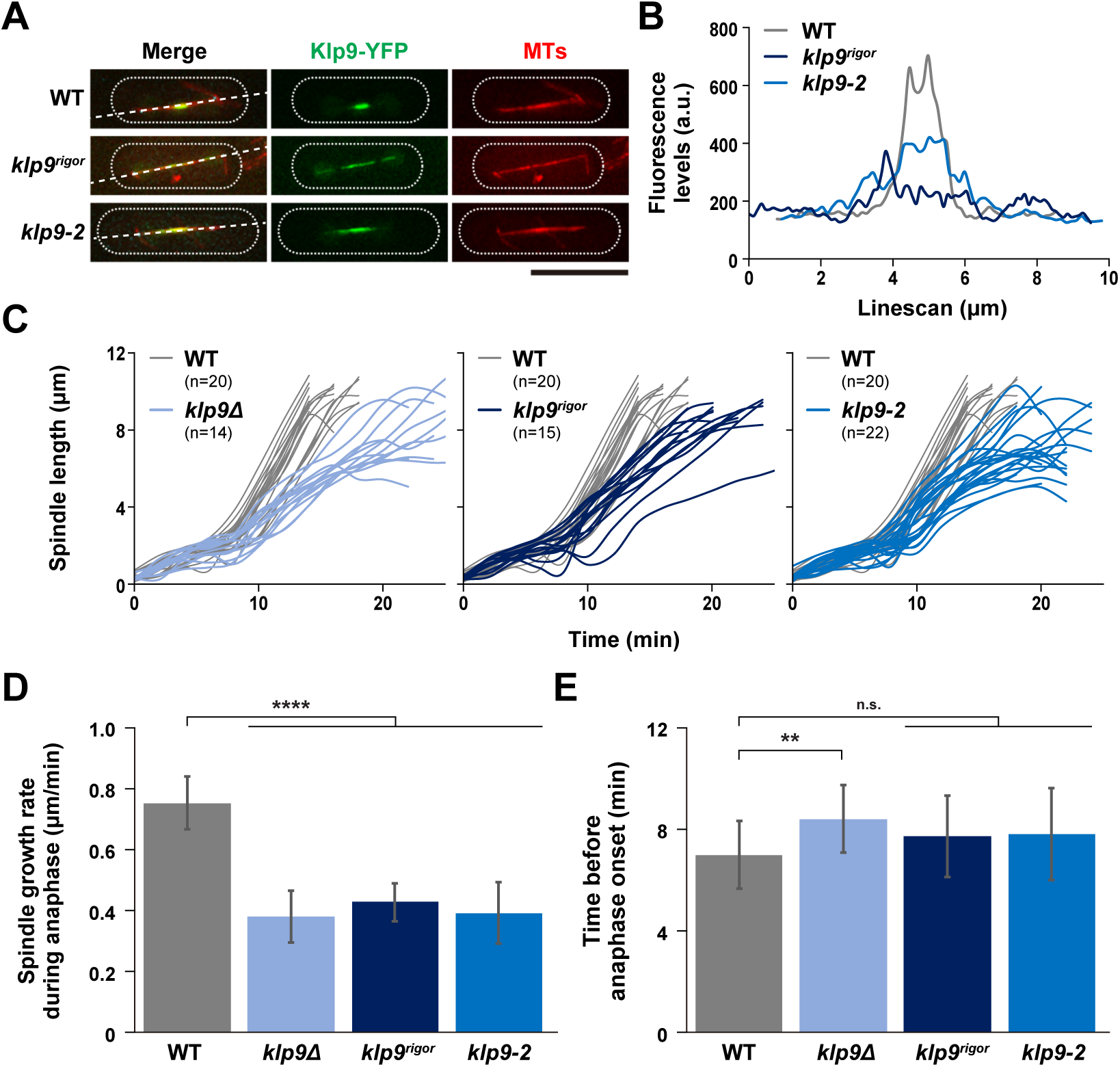
The Klp9 motor is an important determinant for spindle elongation rate during anaphase B. **(A)** Klp9-YFP becomes dispersed along the mitotic spindle in the *klp9* motor mutants. Representative images showing mitotic localisation of Klp9-YFP in the indicated strains are presented. Cells of wild type, *klp9^rigor^* and *klp9-2* (carrying Klp9-YFP and mCherry-Atb2) were shifted from 27°C to 36°C, and incubated for 2 h. Scale bar, 10 µm. Lines used for quantification of Klp9-YFP signals in (**B**) are shown on the left panels. **(B)**Representative line scans of Klp9-YFP in wild type (grey line), *klp9^rigor^* (dark blue line) and *klp9-2* (blue line) taken along the spindle axis and between the two SPBs (dotted lines) as shown in **(A)**. **(C)** Profiles of mitotic progression in *klp9∆* (light blue line, n=14), *klp9^rigor^* (dark blue line, n=15) or *klp9-2* cells (blue line, n=22). Each strain contains a tubulin marker (mCherry-Atb2) and an SPB marker (Cut12-GFP). Cells were grown at 36°C for 4 h and live imaging performed thereafter. Changes of the inter-SPB distance were plotted against time. In each panel, patterns of wild type cells are plotted for comparison (grey line, n=20). **(D)** Spindle growth rate during anaphase B. **(E)** The time between the initiation of SPB separation and onset of anaphase B. Data are given as means ± SD; **, P < 0.01; ****, P < 0.0001; n.s., not significant (two-tailed unpaired Student’s *t*-test).

We then measured spindle elongation rate in these mutants as well as in a *klp9* deletion strain (*klp9Δ*). In all three mutants examined, spindle elongation rate was significantly reduced compared with wild type cells (~50% reduction, 0.75±0.09 for wild type vs 0.38±0.09-0.43±0.06 for the three *klp9* mutants, Fig. 2C and 2D). By contrast, we did not detect any delay regarding anaphase onset in either *klp9^rigor^* or *klp9-2* cells (Fig. 2E). It is of note that, a modest, yet significant delay was observed in *klp9Δ* cells, consistent with a previous report ^45^. This is attributable to a motor-independent role of Klp9 in controlling the timing of anaphase onset (Fig. 2E). Overall, these results indicate that Klp9 is a plus-end motor that is crucial for determining the rate of anaphase B spindle elongation.

### Klp9 plays an additional role during anaphase B independent of its motor activity in collaboration with the microtubule crosslinker Ase1

Ase1 is a homologue of vertebrate PRC1 and plays a vital role in spindle assembly and stability as an antiparallel microtubule bundler/crosslinker ^48-50, 52^. Previous work showed that a *klp9Δase1Δ* double deletion strain is inviable ^37^, which we also found (Fig. 3A top). However, it is not established as to how these two MAPs functionally interact and ensure cell viability in collaboration ^25, 37, 45^. As Klp9 is a plus end-directed motor, the simplest scenario would be that Ase1-mediated microtubule crosslinking and Klp9 motor-dependent microtubule sliding activities are required for spindle integrity in a functionally redundant fashion. To interrogate this possibility, we crossed *ase1Δ* with the motor-defective *klp9-2*. Intriguingly and rather unexpectedly, we found that the resulting *ase1∆klp9-2* double mutant strain was not ts (Fig. 3B). In sharp contrast, *cut7∆pkl1∆* double mutants that require motor activity of Klp9 for survival ^26^ displayed ts lethality in combination with *klp9-2* (Fig. 3B). This result implies that Klp9 motor activity alone does not confer viability of *ase1∆* cells.

**Figure 3.**
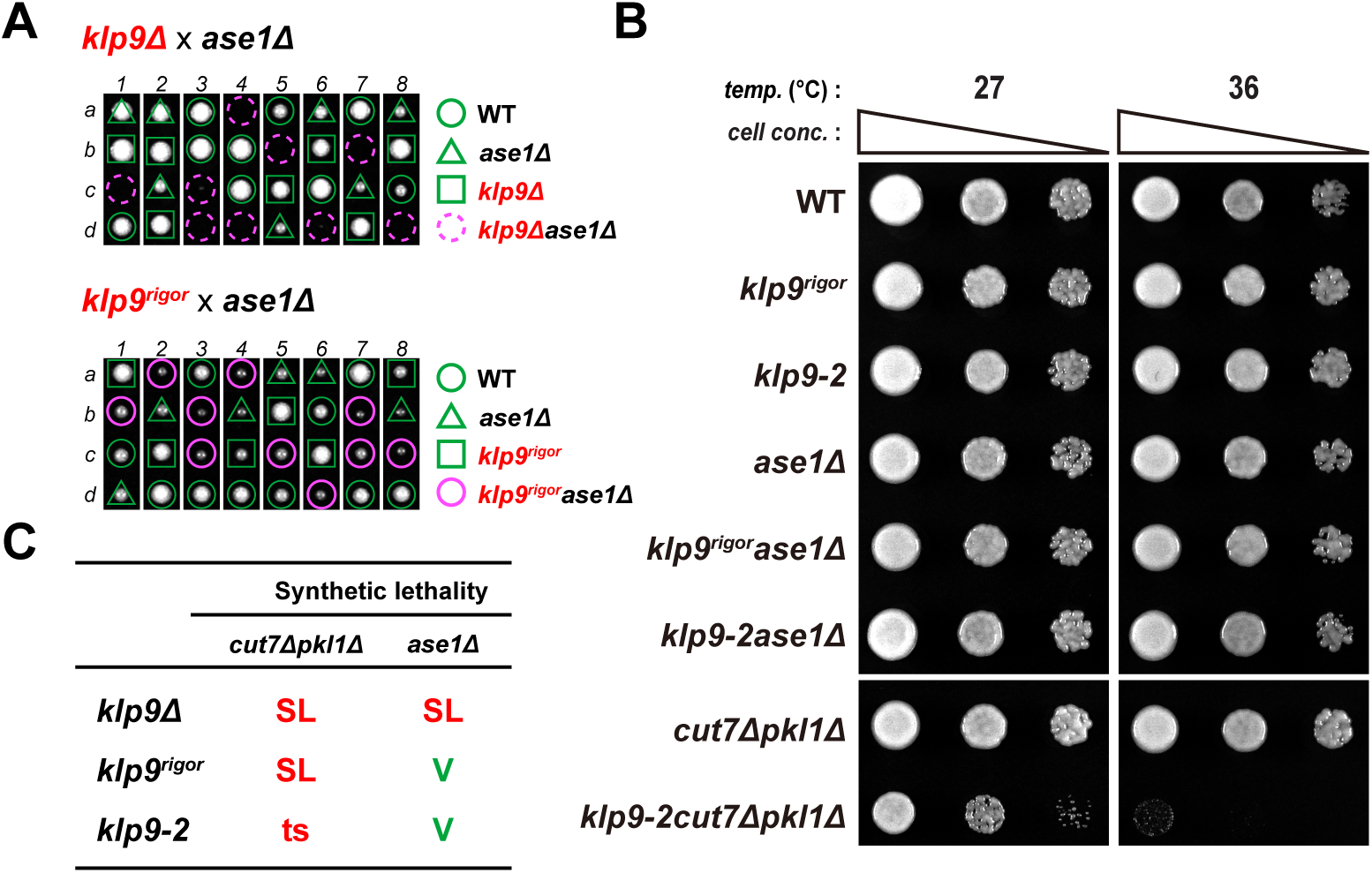
Klp9 plays an additional role during anaphase B independent of its motor activity in collaboration with Ase1. **(A)** Tetrad analysis. Spores were dissected upon crosses between *klp9∆* and *ase1∆* strains (top) or between *klp9^rigor^* and *ase1∆* strains (bottom), respectively. Individual spores (a– d) in each ascus (1–8) were dissected on YE5S plates and incubated for 3 d at 27°C. Representative tetrad patterns are shown; images of dissected tetrads were merged, in which while spaces were created between each tetrad. Circles, triangles and squares with green lines indicate wild type, *ase1∆* single mutants and *klp9* single mutants, respectively. Assuming 2:2 segregation of individual markers allows the identification of lethal *klp9∆ase1∆* double mutants (indicated by dashed magenta circles). Circles with magenta lines indicate lethal *klp9^rigor^ase1∆* double mutants. **(B)** Spot test. Indicated strains were serially (10-fold) diluted, spotted onto rich YE5S plates and incubated at 27°C or 36°C for 2 d. Two parts of the same plate incubated at each temperature were merged, in which while spaces were created between each image. *cell conc.*, cell concentration, *temp.*, temperature. **(C)** Summary of genetic interactions between the *klp9* mutants (*klp9∆*, *klp9^rigor^* or *klp9-2*) and *cut7∆pkl1∆* or between the *klp9* mutants and *ase1∆*. SL, synthetic lethal. V, viable. ts, temperature sensitive.

To substantiate this notion, we crossed *klp9^rigor^* with *ase1Δ* and found that the double mutant was indeed viable and capable of forming colonies at any temperatures including 36°C (Fig. 3A, bottom and 3B). This indicates that defective motor activity of Klp9 is not responsible for synthetic lethality between *ase1∆* and *klp9∆*. In order to decipher the essential role of Klp9 in the absence of Ase1, we next investigated the functional domain within Klp9 that is required for viability of *ase1Δ* cells.

### Systematic truncation and mutation analysis of Klp9 highlights the importance of an NLS in the C-terminal non-motor region

It was reported that Klp9 plays one additional, motor-independent role in regulating the timing of anaphase onset (see Fig. 2E), which requires the C-terminal 38 amino acid residues ^45^. Crossing *ase1∆* with the *klp9* mutant lacking this region (*klp9-∆38C*) showed that the double mutant was still viable (Fig. 4A and Supplementary Fig. 2), indicating that the role of Klp9 in anaphase onset is not involved in the functional redundancy with Ase1. We then systematically created a series of C-terminally truncated *klp9* mutants at the endogenous locus under the native promotor with GFP in the C-terminus, and crossed them with *ase1∆*. These included Klp9-∆92C, Klp9-∆133C and Klp9-∆172C, which lacks one of the two coiled coil domains and Klp9-∆234C that does not contain either of coiled coil domains (Fig. 4A) and finally, Klp9-∆Motor that lacks the entire N-terminal motor domain. It was found that all the truncation mutants except for *klp9-∆38C* were synthetically lethal with *ase1∆* (Fig. 4A and Supplementary Fig. 2).

**Figure 4.**
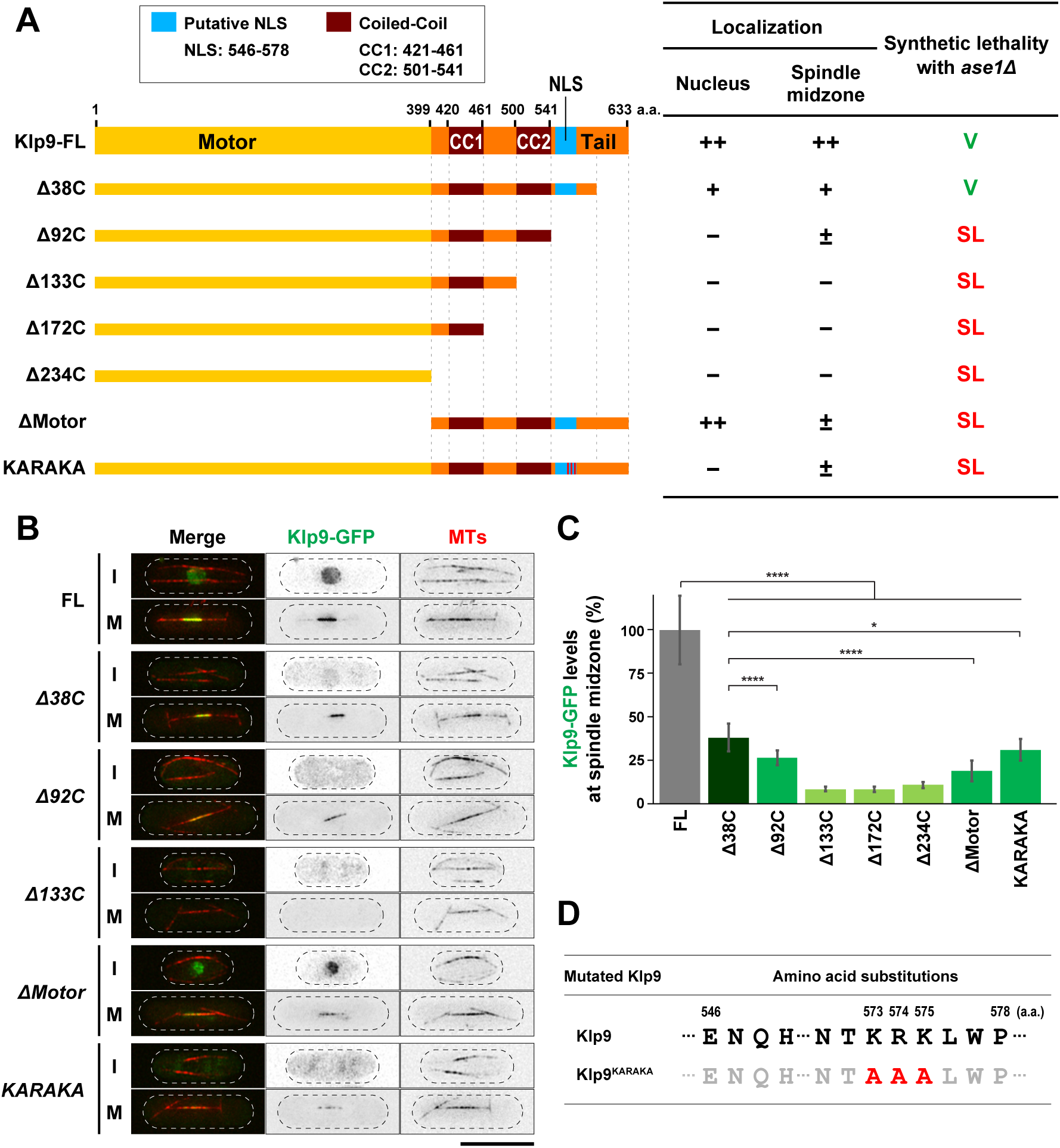
Systematic truncation and mutation analysis of Klp9 highlights the importance of NLS in the C-terminal non-motor region. **(A)** Schematic representation of full-length Klp9 (Klp9-FL), C-terminal truncation mutants (***∆***38C, ***∆***92C, ***∆***133C, ***∆***172C and ***∆***234C), an NLS mutant (KARAKA) and N-terminal truncation mutant (***∆***Motor). The table on the right indicates a summary of the cellular localisation and the synthetic lethality of each *klp9* mutant. ++, +, ± and − indicate normal, low, very low and invisible levels of Klp9-GFP localisation in the nucleus or the spindle midzone, respectively. SL, synthetic lethal. V, viable. **(B)** Representative images showing Klp9-GFP localisation during interphase (I) and mitosis (M) in the indicated strains are presented. Scale bar, 10 µm. **(C)** Quantification of Klp9-GFP levels at the midzone of spindle microtubules during anaphase B. Data are given as means ± SD; *, P < 0.05; ****, P < 0.0001 (two-tailed unpaired Student’s *t*-test). (**D**) Substituted amino acid residues in *klp9^KARAKA^* mutant. These point mutations (K573AR574AK575A) were introduced upon site-directed mutagenesis and confirmed by nucleotide sequencing of the *klp9* gene on the selected clone.

We also observed the cellular localisation of individual Klp9 mutants during interphase and mitosis. Previous work showed that full-length Klp9 is localised to the nucleus throughout the cell cycle, but its localisation pattern differs depending upon cell cycle stages; it occupies the entire nucleoplasm from interphase until mid-mitosis, and then translocates to the spindle midzone in late mitosis ^37^. First, we confirmed this profile (Fig. 4A and 4B, top row). Interestingly, Klp9-∆92C, but not Klp9-∆38C or Klp9-∆Motor, lost nucleoplasmic localisation during interphase. We noted that the mitotic localisation of Klp9-∆92C to the spindle midzone was somehow still retained, though precise quantification of signal intensities showed a substantial reduction compared to those in wild type cells (Fig. 4B and 4C). By contrast, all the other truncation mutants (Klp9-∆133C, -∆172C, -∆234C) did not localise to either the nucleus or the spindle midzone (Fig. 4A-4C).

Nuclear localisation of Klp9 during interphase suggested the presence of a nuclear localisation signal (NLS) ^53^. Computational search picked up a putative NLS between 546^th^ and 578^th^ that contains three consecutive basic amino acids (573K-574R-575K, Fig. 4D). The replacement of three basic residues with alanines (designated Klp9-KARAKA) abolished the nuclear localisation during interphase despite that Klp9-KARAKA was capable of weakly localising to the anaphase B spindle like Klp9-∆92C (Fig. 4A-4C). Importantly, *klp9-KARAKA* was synthetically lethal with *ase1∆* (Fig. 4A, 4B and Supplementary Fig. 2); thus *klp9-KARAKA* recapitulated defective profiles of *klp9-∆92C*. It should be noted that all the mutants created, except for *klp9-∆38C*, also exhibited synthetic lethality in combination with *cut7∆pkl1∆* (Supplementary Fig. 2). Collectively, the domain analysis shown here has unveiled the existence of the NLS that is essential for correct localisation and function of Klp9.

### Two internal coiled coil regions are crucial for Klp9 function in collaboration with Ase1

Next, we created internal deletion mutants of Klp9. The C-terminal region of Klp9 contains two coiled coil domains (CC1, 421-461 and CC2, 501-541). We deleted each region and, integrated it into the endogenous locus with GFP in the C-terminus (designated Klp9-∆CC1 and Klp9-∆CC2 respectively) (Fig. 5A). The cellular localisation of these Klp9 mutants showed that neither of them was localised to the spindle midzone in late mitosis (Fig. 5B and 5C). It is noteworthy that interphase nuclear localisation was retained in these mutants, consistent with the notion that they contain the C-terminal NLS identified earlier (Fig. 4). Genetic analysis indicated that both deletion mutants were synthetically lethal with either *ase1∆* or *cut7∆pkl1∆* (Fig. 5D and Supplementary Fig. 2), implying that both CC1 and CC2 are indispensable for Klp9 function. These result highlighted that the C-terminal coiled coil domains are essential for Klp-9’s plus end-directed motor activity as well as its motor-independent role in collaborating with Ase1.

**Figure 5.**
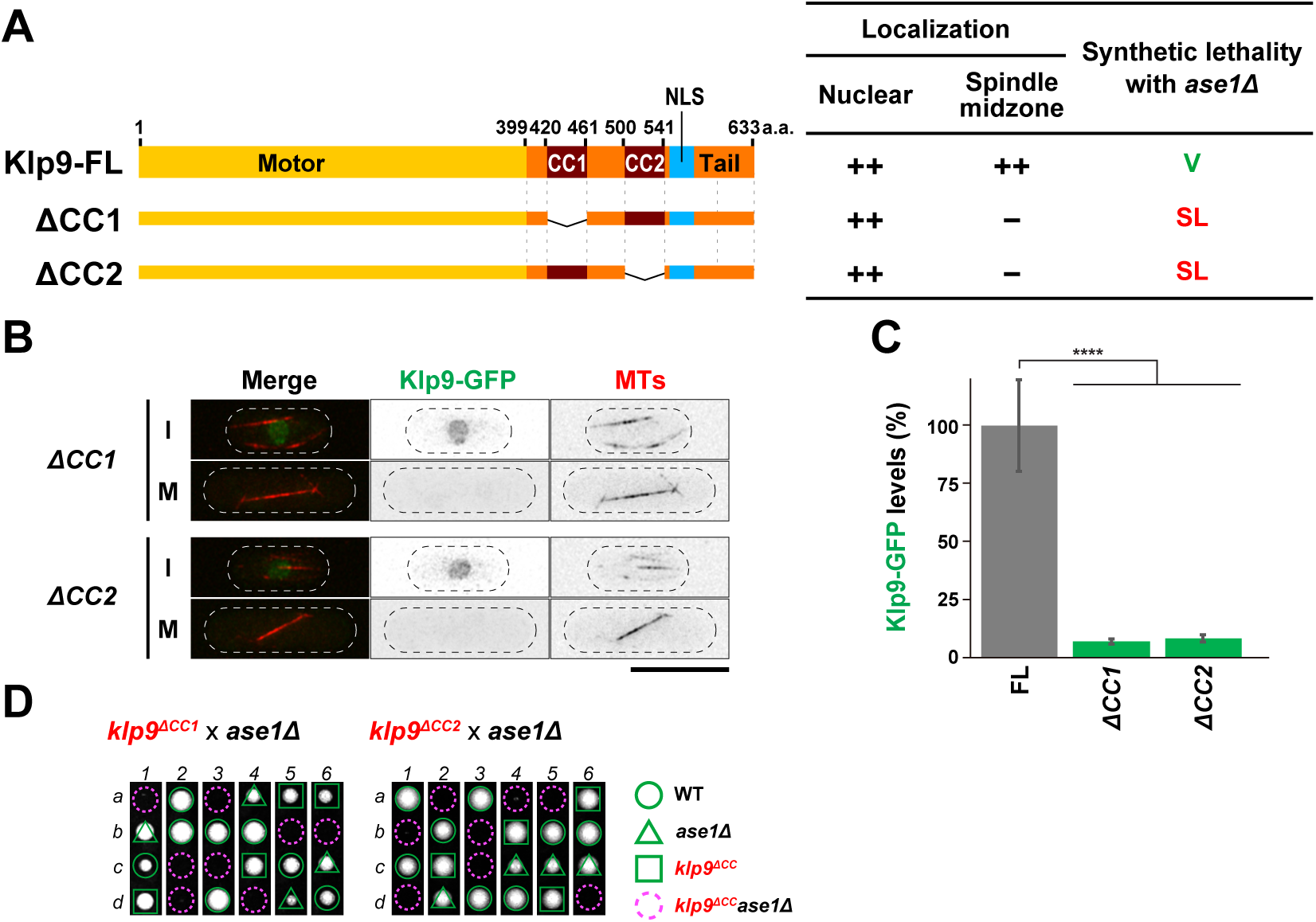
Two internal coiled coil regions are crucial for Klp9 function in collaboration with Ase1. **(A)**Schematic representation of full-length Klp9 (Klp9-FL) and two internal deletion mutants. Each of coiled coil domains (Klp9-∆CC1: ∆421-461 and Klp9-∆CC2: ∆501-541) is deleted. The table on the right indicates a summary of the cellular localisation and the synthetic lethality of each *klp9* mutant. ++ and − indicate normal and invisible levels of Klp9-GFP localisation in the nucleus or the spindle midzone, respectively. SL, synthetic lethal. V, viable. **(B)** Representative images showing Klp9-GFP localisation in the indicated strains during interphase (I) and mitosis (M) are presented. Scale bar, 10 µm. **(C)** Quantification of Klp9-GFP levels at the midzone of spindle microtubules during anaphase B. Data are given as means ± SD; ****, P < 0.0001 (two-tailed unpaired Student’s *t*-test). **(D)** Tetrad analysis. Spores were dissected upon crosses between *klp9^∆CC1^* and *ase1∆* strains or between *klp9^∆CC2^* and *ase1∆* strains respectively. Individual spores (a–d) in each ascus (1–6) were dissected on YE5S plates and incubated for 3 d at 27°C. Representative tetrad patterns are shown; images of dissected tetrads were merged, in which while spaces were created between each tetrad. Circles, triangles and squares with green lines indicate wild type, *ase1∆* single mutants and *klp9^∆CC^* single mutants, respectively. Assuming 2:2 segregation of individual markers allows the identification of lethal *klp9^∆CC^ase1∆* double mutants (indicated by dashed magenta circles).

### Kinesin-5 Cut7 collaborates with Klp9 in spindle elongation during anaphase B

In *klp9* mutants with defects in motor activity (eg. Klp9-2 and Klp9^rigor^), spindle elongation during anaphase B was substantially compromised; however, spindles were still capable of elongating, albeit with a reduced rate (see Fig. 2C and 2D). Previous studies suggested the contribution of Cut7 in late mitosis ^25, 26, 54^. However, there is no direct evidence for the requirement of Cut7 in anaphase B spindle elongation and this issue remains under debate ^37^. One of the reasons for this conundrum is that Cut7 is required in the early stages of spindle assembly. As defects in SPB separation leads to the formation of monopolar spindles at restrictive temperatures, all available *cut7* ts mutants do not reach anaphase B ^5, 18, 54^. We posited that if Cut7 is also involved in a later step of spindle assembly, a novel type of conditional mutants could be isolated, in which they are defective only during late mitosis and exhibit the ts phenotype only in the background of *klp9∆*; in other words, such *cut7* mutants would not be ts for growth on their own.

Given this assumption, we introduced PCR-mutagenised *cut7* fragments into *klp9∆* cells and screened for ts transformants (see Methods for details). We then backcrossed individual ts isolates with wild type cells and examined how the ts phenotype segregated in progenies; expected *cut7* mutants should only display temperature sensitivity in the background of *klp9∆*. A number of anticipated ts mutants were successfully identified (details will be reported elsewhere). In this study, we present one representative mutant called *cut7-122*. As shown in Fig. 6A, *cut7-122* cells displayed the ts phenotype only in the *klp9∆* background, but not in the presence of Klp9. Nucleotide sequencing of the *cut7-122 ORF* identified two point mutations, which led to amino acid replacements at 134^th^ and 290^th^ residues (M134T and T290K) within the N-terminal motor domain (Fig. 6B).

**Figure 6.**
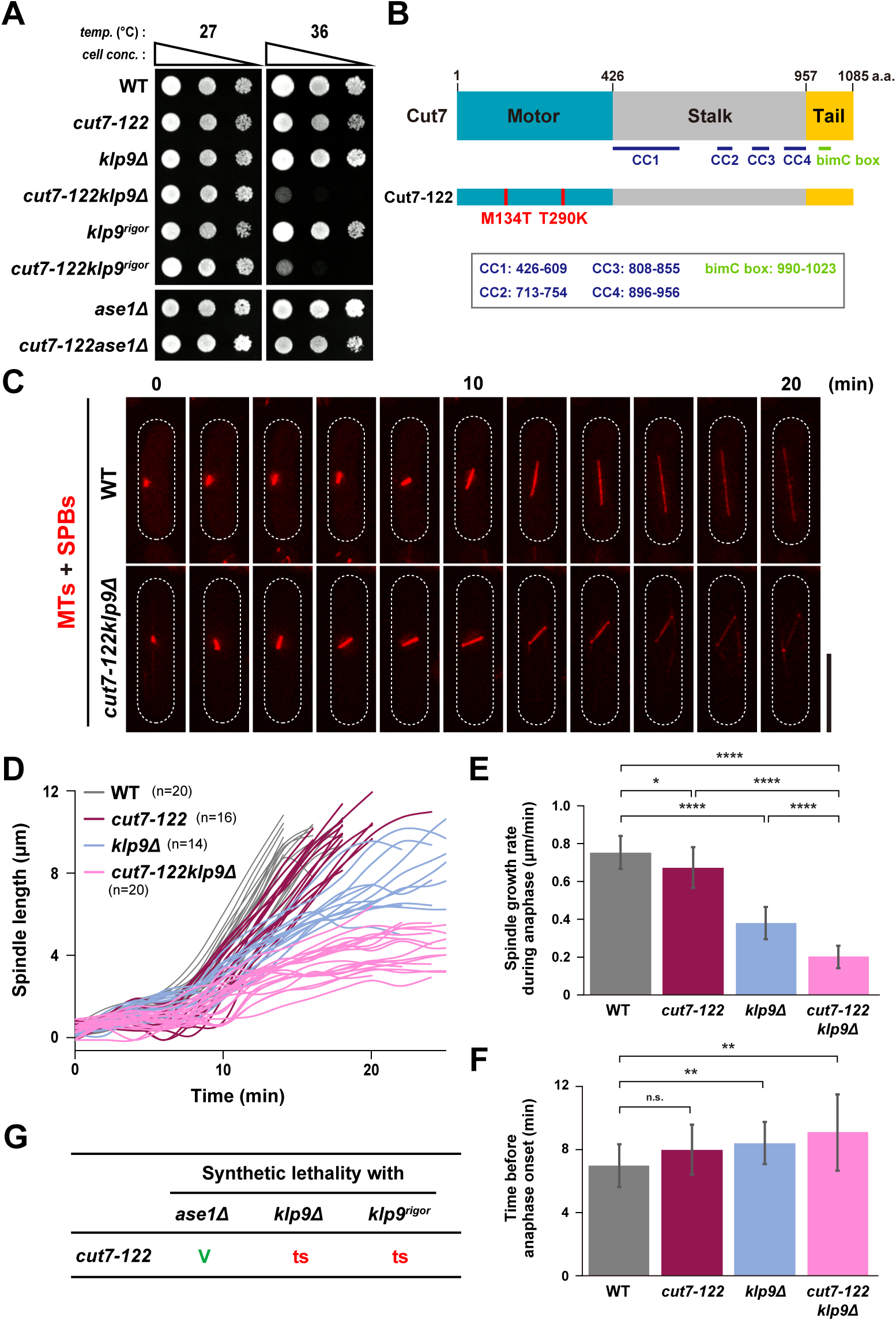
Kinesin-5 Cut7 collaborates with Klp9 in spindle elongation during anaphase B. **(A)** Spot test. Indicated strains were serially (10-fold) diluted, spotted onto rich YE5S plates and incubated at 27°C or 36°C for 2 d. *cell conc.*, cell concentration, *temp.*, temperature. Two parts of the same plate incubated at each temperature were merged, in which while spaces were created between each image. **(B)** Mutation sites in the *cut7-122* mutant. Cut7-122 contained two mutations (M134T and T290K) in the C-terminal motor domain. **(C)** Time-lapse images of mitotic wild type and *cut7-122klp9Δ* cells. Spindle microtubules (mCherry-Atb2; red) were visualised. Images were taken at 2 min intervals after incubation of cultures at 36°C for 4 h. The cell peripheries are outlined with dotted lines. Scale bar, 10 µm. **(D)** Profiles of mitotic progression in *cut7-122* (dark magenta line, n=16) or *cut7-122klp9Δ* cells (light magenta line, n=20). Each strain contains a tubulin marker (mCherry-Atb2). Cells were grown at 36°C for 4 h and live imaging performed thereafter. Changes of the inter-SPB distance were plotted against time. Data for wild type (grey line, n=20) and *klp9Δ* (light blue line, n=14) are plotted for comparison and these are the same as those presented in Fig. 2C. **(E)** Spindle growth rate during anaphase B. **(F)** The time between the initiation of SPB separation and onset of anaphase B. Data for wild type and *klp9Δ* are shown for comparison; these are the same as those presented in Fig. 2D and 2E. Data are given as means ± SD; *, P < 0.05; **, P < 0.01; ****, P < 0.0001; n.s., not significant (two-tailed unpaired Student’s *t*-test). **(G)** Summary of genetic interactions between *cut7-122* and *klp9* mutants or *ase1∆.* V, viable. ts, temperature sensitive.

To study the impact of the *cut7-122* mutant on mitotic spindle elongation on its own or in combination with *klp9∆*, we observed the dynamic behaviour of spindle microtubules in *cut7-122* or *cut7-122klp9∆* cells expressing mCherry-Atb2 (a microtubule marker) ^51^ and Sid4-mRFP (an SPB marker) ^55^. Intriguingly, elongation rate of anaphase B spindles was decreased in a single *cut7-122* mutant: 0.67±0.11 μm/min and 0.75±0.09 μm/min for *cut7-122* and wild type cells respectively. Furthermore, *cut7-122klp9∆* double mutants displayed a very low rate of spindle elongation (0.20±0.06 μm/min, Fig. 6C-6E); Cut7 and Klp9 contributed additively to the elongation of anaphase B spindles. It is of note that a single *cut7-122* mutant did not display a significant delay in anaphase onset (Fig. 6F) unlike previously isolated conventional *cut7* alleles ^5, 25, 26, 56^. This sustains the notion that the major, if not sole, defect in this novel *cut7* allele lies in the late stage of mitosis.

Given the synthetic temperature sensitivity between *cut7-122* and *klp9∆*, we next asked which activity of Klp9 is required for this genetic interaction. We found that *cut7-122* displays the ts phenotype in combination with *klp9^rigor^*, which is in stark contrast to the result that *cut7-122* did not exhibit the ts growth in combination with *ase1∆* (Fig.6A and 6G). Therefore, the survival of *cut7-122* depends upon the motor activity of Klp9. The profiles of genetic interactions obtained for *klp9^rigorI^/klp9-2* and *cut7-122* are identical with regards to Ase1 (Figs. 3C and 6G). Thus, we conclude that Cut7 functions in late mitosis; it acts in concert with the motor activity of Klp9 in spindle elongation during anaphase B.

## DISCUSSION

In this study, we explored the molecular mechanisms by which mitotic spindles elongate during anaphase B in fission yeast. Previous work showed the importance of Kinesin-6 Klp9 and the microtubule crosslinker Ase1 ^26, 37, 45^; however the molecular details underlying the functional interplay between these two MAPs are not known. In addition, it is undetermined whether Kinesin-5 Cut7 plays any role in late mitosis. Using biophysical, cell biological and genetic approaches, we show that Klp9 is a processive motor with plus-end directionality and collaborates with both Ase1 and Cut7 during anaphase B, in two distinct manners. The former collaboration is independent of Klp9 motor activity, while the latter role relies on outward force generated by Klp9 as a motor. Furthermore, isolation of a novel *cut7* mutant allele that is defective specifically in anaphase B spindle elongation has solidified the notion that the Cut7 motor is indeed required for spindle elongation, not only in an early mitotic stage, but also a later step as well (a model illustrated in Fig. 7).

**Figure 7.**
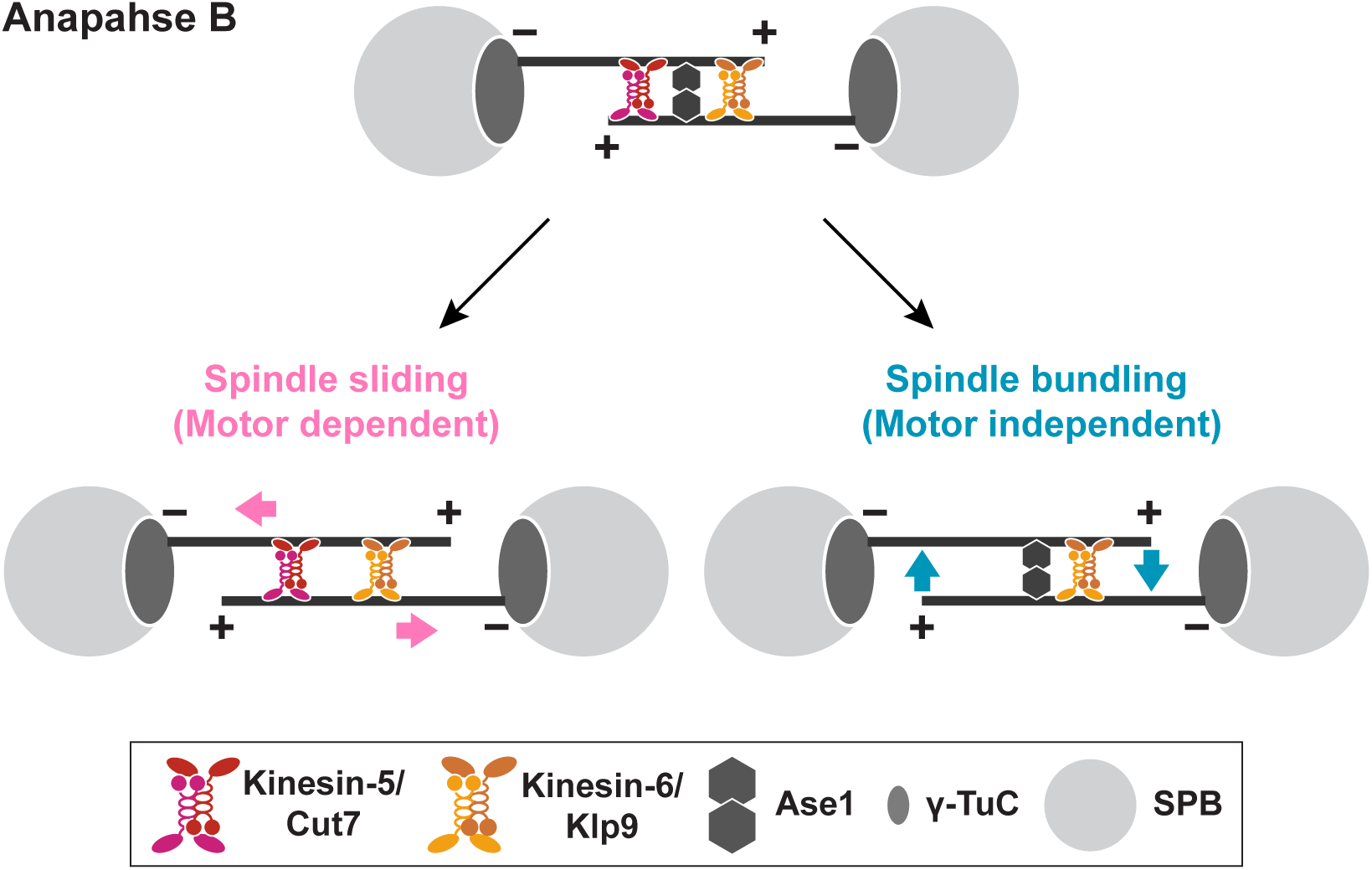
A model of spindle elongation during anaphase B. Spindle elongation during late mitosis is executed by three MAPs, Kinesin-6 Klp9, Kinesin-5 Cut7 and the microtubule crosslinker Ase1. Klp9 plays dual roles in this process; one is motor-dependent and the other is motor-independent. Klp9 motor activity generates outward force, thereby inducing the sliding away of antiparallel microtubules at the spindle midzone (bottom left). Cut7 also generates outward force in an additive manner. This force-generating process collaborates with Ase1 to maintain microtubule bundling of anaphase B spindles and that Klp9 plays a role in this branch independently of its motility (bottom right). Motor-independent role of Klp9 requires the NLS and two coiled coil domains locating at the C-terminal non-motor region. Cut7 may not be involved in this microtubule crosslinking; or at least its role is less important compared with that played by Klp9.

### Kinesin-6 Klp9 is a processive plus end-directed motor

Our in vitro TIRFM assay explicitly shows that Klp9 moves towards microtubule plus ends in a processive manner (Fig.1). The measured velocity of 21 nm/s is comparable with other reported values for plus end-directed Kinesin-5 motors ^57–59^.

Recently, it has become clear that fungal Kinesin-5s including Cut7 are bidirectional motors; they display either plus end-directed or minus end-directed motilities depending upon conditions (eg. the degree of motor crowding on the microtubule lattice) ^59–61^. Under our experimental conditions, we did not detect a similar bi-directionality, despite Klp9 forming homotetramers like Cut7. It would be of interest to address where this difference stems from and ask for the physiologies behind their distinct properties.

### Dual roles of Kinesin-6 Klp9 in spindle elongation and microtubule bundling during anaphase B

We show that Klp9 does not require its motor activity to carry out essential functions in the absence of Ase1 (Fig. 3). Instead, it depends upon the two domains within the C-terminal non-motor region: two coiled coils and the NLS (Figs. 4 and 5). It is shown that Klp9 forms homotetramers like Cut7 ^37^ and that the analogous C-terminal coiled coils in Cut7 and other Kinesin-5 members are responsible for self-tetramerisation ^62–64^. We envision that coiled coil-dependent homotetramers of Klp9 play a structural role in the bundling of antiparallel microtubules at the spindle midzone together with Ase1. To our knowledge, this is the first report on kinesin molecules that play a motor-independent role in microtubule crosslinking in a physiological context. Whether Kinesin-6 members in other species play an analogous role would be of great interest to be explored.

### Roles of Kinesin-5 Cut7 during late mitosis

Using a newly isolated *cut7* ts allele, *cut7-122*, our analysis uncovered the crucial role of Cut7 as a motor in late mitosis. Spindle elongation rate in a single *cut7-122* mutant is reduced by 11% compared to wild type cells, while that in a *klp9∆* strain drops by 49% and in a *cut7-122klp9∆* double mutant, the reduction reaches 73% (Fig. 6E). As the *cut7-122* strain is most likely hypomorphic, not a complete loss of function, this degree of contribution from Cut7 to anaphase B spindle elongation is potentially underestimated.

To address how these two motors collaborate with each other on the antiparallel microtubules, performing TIRFM ^65^ in the presence of individual proteins or a mixture of the two as a microtubule pair sliding assay would provide useful information.

Despite its slow velocity, we still detect spindle elongation in the *cut7-122klp9∆* double mutant (retaining 27% activity compared to wild type cells). As aforementioned, although the Cut7-122 proteins are unlikely to have completely abolished motor activity, cells might still be capable of undergoing spindle elongation without kinesin-generated motor activities. This could be ascribable to the outward forces elicited by non-motor MAPs, including microtubule polymerisation and/or Ase1- and CLASP/Cls1-mediated microtubule bundling and stabilisation as previously pointed out ^25, 26^.

Although the requirement of Cut7 for anaphase B spindle elongation has been established in this study, whether Cut7 plays another motor-independent role in microtubule bundling, analogous to Klp9, remains undetermined. We observed that *klp9* mutants which cannot form homotetramers are synthetically lethal with *ase1∆* regardless of Cut7’s presence (Fig. 5A). This raises the possibility that Cut7 may not be involved in microtubule crosslinking or at least its role is not as critical as that of Klp9. Thus, activities and/or properties of microtubule binding may not be identical between Cut7 and Klp9. Further cell biological and in vitro assays are required to clarify whether Cut7 is implicated in spindle integrity during anaphase B in a motor-independent manner.

### Cell cycle-dependent cellular localisation of Kinesin-6 Klp9

We have identified a canonical NLS in the C-terminal region of Klp9 that is essential for its mitotic function. Klp9 constructs without a functional NLS could still be loaded onto the spindle midzone within the nucleus, though with reduced levels (Fig. 4A and 4B). Thus, it is likely that Klp9 is transported into the nucleus in a dual manner: one through its own NLS and the other perhaps through binding to another protein. Ase1, which acts together with Klp9 during anaphase B, might well be such a binding partner; however, two previous reports ^37, 45^ as to whether Klp9 physically binds Ase1 are somewhat controversial. Hence, how Klp9 is localised to the mitotic nucleus in an NLS-independent manner remains undetermined. However, it is worth noting that despite the capability of its NLS-independent nuclear import during late mitosis, levels of Klp9 on spindles are insufficient to support the mitotic role of Klp9 (Fig. 4A). Thus, the NLS within the C-terminal domain of Klp9 is essential for Klp9 function.

### Kinesin-5 and Kinesin-6 in spindle elongation in other species

The role of Kinesin-5 in bipolar spindle assembly, in particular during very early mitotic stages when the two centrosomes separate, is conserved across many eukaryotic species ^66^. In addition, Kinesin-6 members ubiquitously participate in late mitotic events. Yet, the functional interplay between these two kinesins, if any, remains to be reported ^10^. The results presented in the current study are the first to explore this notion. In budding yeast, Kinesin-5 Cin8 concentrates on central spindle microtubules during anaphase and plays a critical role in the maintenance of spindle pole separation ^8, 67^. However, this organism does not contain Kinesin-6 members. Instead, it has another Kinesin-5 Kip1, which collaborates with Cin8 in spindle elongation. It is likely that this yeast has developed a unique evolutionary strategy by which two different Kinesin-5 members compensate for the functions of Kinesin-6 in anaphase B spindle elongation. It is of note that a recent study showed that Kinesin-8 Kip3 acts in late mitosis together with Kip1 ^68^.

In higher eukaryotes, multiple Kinesin-6 proteins play collaborative roles in spindle elongation during anaphase B coupled with cytokinesis. In *Drosophila melanogaster*, Kinesin-5 Klp61F and Kinesin-6 Subito work together in maintaining meiotic bipolar spindles ^69^. Subito is known to be an important kinesin for mitotic spindle assembly as well. It would be of great interest to address whether Klp61F and Subito’s collaboration in spindle assembly extends into mitosis.

## MATERIALS AND METHODS

### Strains, media, and genetic methods

Fission yeast strains used in this study are listed in Supplementary Table S1. Media, growth conditions, and manipulations were carried out as previously described ^70–72^. For most of the experiments, rich YE5S liquid media and agar plates were used. Spot tests were performed by spotting 5–10 µl of cells at a concentration of 2 × 10^7^ cells/ml after 10-fold serial dilutions onto rich YE5S plates. Plates were incubated at various temperatures from 27°C to 36°C as necessary.

### Preparation and manipulation of nucleic acids

Enzymes were used as recommended by the suppliers (New England Biolabs Inc. Ipswich, MA, U. S. A. and Takara Bio Inc., Shiga, Japan).

### Construction of recombinant Klp9 for transient expression

To construct vectors for transient expression of recombinant Klp9 (pcDNA3.4/SpKlp9(full)-EGFP-FLAG-His8), the full-length *klp9* cDNA was PCR amplified from a cDNA library (National BioResource Project, pTN-RC5) with a forward primer, ATGATACAGATTTTTCTGCGTGT and a reverse primer, AATTCATTAATATCGATATCAGTTG, and inserted into a modified pcDNA3.4 vector (pcDNA3.4/EGFP-FLAG-His8) via appropriate oligo nucleotide adaptors for the acceptor vector using the Gibson Assembly method. The resultant linker amino acid between SpKlp9 and EGFP is AAA (one-letter amino-acid code).

### Expression and purification of Klp9

We essentially followed procedures described previously ^73^. For transient expression, Expi293F cells (A14635, Invitrogen) were used according to the manufacturer's instructions except that transfection was performed with PEI ‘Max’ (24765-2, Polysciences). Cells from 800 ml culture were collected by centrifugation at 1,930 r.p.m. (700×g) for 10 min, washed three times with PBS buffer (pH 7.4), and stored at −80° C. Frozen cells were suspended in lysis buffer (20 mM Tris-HCl at pH 7.5, 250 mM NaCl, 1 mM MgSO_4_, 100 μM ATP, 10% (w/v) sucrose, 10 mM imidazole-HCl at pH 7.5 and 0.5 mM dithiothreitol) containing 1 mM phenylmethylsulphonyl fluoride and a protease inhibitor cocktail (03969-21, Nacalai Tesque). The resuspended cells were then disrupted by a sonicator (Sonifier 250, Branson). After ultracentrifugation at 48,000 r.p.m. (21,000×g) for 45 min at 4°C, the supernatant was loaded to Ni-IMAC resin (156-0133, Bio-Rad) and eluted with 20 mM Tris-HCl at pH 7.5, 250 mM NaCl, 1 mM MgSO_4_, 100 µM ATP, 10% (w/v) sucrose, 250 mM imidazole-HCl at pH 7.5 and 0.5 mM dithiothreitol. The eluate was next loaded to anti-FLAG agarose (A2220, Sigma-Aldrich) and eluted with lysis buffer supplemented with 350 µg/ml 3×FLAG peptide (F4799, Sigma-Aldrich). The eluate was flash frozen in liquid nitrogen and stored at –80°C. Protein concentration was determined by the Bradford assay using bovine serum albumin as a standard.

### Preparation of fluorescently labelled and polarity-marked microtubules

Tubulin was purified from porcine brain using a high-molarity PIPES buffer (1 M PIPES-KOH pH6.8, 20 mM EGTA, and 10 mM MgCl_2_) as described previously ^74^. To prepare fluorescently labelled microtubules, tubulin was labelled with Cy3 (PA23001, GE Healthcare) or ATTO 647N (AD 647N-31, ATTO-TEC). ATTO 647N-microtubules were polymerised by copolymerising labelled tubulin and unlabelled tubulin at a ratio of 1:5 for 30 min at 37°C, and stabilised with 40 µM paclitaxel (T1912, Sigma-Aldrich). Polarity-marked microtubules were prepared as described previously ^73^.

### Microtubule gliding assay

A flow chamber was first filled with 7 µl of antibody to penta-His (#34660, Qiagen; 1/10 dilution) and incubated for 5 min. The chamber was next coated with 1%(w/v) Pluronic F-127 (P2443, Sigma-Aldrich), allowed to adsorb for 5 min and blocked with ~0.7 mg/ml casein (#07319-82, Nacalai Tesque) in BRB80 buffer (80 mM PIPES-KOH pH6.8, 1 mM EGTA and 1 mM MgCl_2_). The flow chamber was then incubated with 40 µg/ml Klp9 solution in BRB80 buffer for 5 min. After washing with 21 µl of BRB80 buffer containing the fluorescently labelled polarity-marked microtubules, 0.218 mg/ml glucose oxidase (G2133, Sigma-Aldrich), 0.04 mg/ml catalase (219001, Calbiochem), 25 mM glucose, ~0.7 mg/ml casein, 2 mM DTT, 10 µM paclitaxel and 1 mM ATP in BRB80 buffer, was introduced into the flow chamber. After a 15 s incubation, final solution without microtubules was perfused into the flow chamber. Microtubule gliding was imaged using an objective-type TIEFM based on Ti-E (Nikon) equipped with a ×60/NA1.49 oil immersion objective lens (CFI Apo TIRF 60xH, Nikon) at 24 ± 1°C. Images were magnified by 2.5× TV-adaptor (MQD42120, Nikon) and projected onto an EMCCD detector (C9100-13, Hamamatsu Photonics). The camera was controlled by the Micro-manager software ver. 1.4.2243. The position of the filament was determined manually by mouse clicking on the tip of the filament in the standalone custom software (Mark2) and tracked typically every 10 s. Velocities of microtubules that were translocated for at least 10 s were calculated by the linear fit to each trace. The stationary microtubules were excluded from the velocity measurement.

### Single molecule motility assay

To immobilise microtubules, the flow chamber was first coated with 10 µg/ml anti-tubulin antibody (SC-58884, Santa Cruz) in BRB80 buffer, allowed to adsorb for 5 min and blocked with 1% (w/v) Pluronic F-127 in BRB80 buffer. After washing with 0.7 mg/ml casein in BRB80 buffer, the flow chamber was incubated with ATTO 647N-labelled microtubules in BRB80 buffer for 5 min. After washing with casein solution, the chamber was filled with final solution containing 0.3 nM Klp9-EGFP, 12 mM PIPES-KOH pH 6.8, 2 mM MgSO_4_, 1 mM EGTA, 10 mM K-acetate, 10 µM paclitaxel, 0.7 mg/ml casein, 2 mM dithiothreitol, 25 mM glucose, 21.3 U/ml glucose oxidase, 800 U/ml catalase and 1 mM ATP. To determine the polarity of the microtubules, the chamber was perfused with kinesin labelled with Alexa546 after measurement. Images were recorded at 300 ms per frame. The positions of the fluorescent spots in each frame were determined by two-dimensional Gaussian fitting with custom software (Mark2). The fluorescent spots that stayed on microtubules for over 4 frames were traced. Velocities were calculated by a linear least-squares fit to each trace.

### Strain construction, gene disruption and the N-terminal and C-terminal epitope tagging

A PCR-based gene-targeting method ^71, 72^ was used for complete gene disruption and epitope tagging in the C terminus, by which all the tagged proteins were produced under the endogenous promoter.

### Isolation of *cut7* temperature-sensitive mutants in the *klp9∆* background

The *cut7* ts mutants were constructed by PCR-based random mutagenesis. The GFP epitope tag and G418-resistance marker gene cassette (*kanR*) were first inserted into the 3’ flanking region of the *cut7* gene (*cut7-GFP-kanR*). The *cut7-GFP-kanR* fragment purified from this strain was amplified/mutagenised with PCR using *TaKaRa EX* taq polymerase (Takara Bio Inc. Shiga, Japan) and transformed into a *klp9∆* strain (deleted by *hphR*). G418-(and Hygromycin B)-resistant colonies were picked up and backcrossed with a wild type strain. After confirming co-segregation between G418/Hygromycin B-resistance and the ts phenotype, nucleotide sequencing was performed to determine the mutated sites within the *cut7* gene of these mutants.

### Fluorescence microscopy and time-lapse live cell imaging

Fluorescence microscopy images were obtained by using a DeltaVision microscope system (DeltaVision Elite; GE Healthcare, Chicago, IL) comprising a wide-field inverted epifluorescence microscope (IX71; Olympus, Tokyo, Japan) and a Plan Apochromat 60×, NA 1.42, oil immersion objective (PLAPON 60×O; Olympus Tokyo, Japan). DeltaVision image acquisition software (softWoRx 6.5.2; GE Healthcare, Chicago, IL) equipped with a charge-coupled device camera (CoolSNAP HQ2; Photometrics, Tucson, AZ) was used. Live cells were imaged in a glass-bottomed culture dish (MatTek Corporation, Ashland, MA) coated with soybean lectin and incubated at 27°C for most of the strains or at 36°C for the ts mutants. The latter were cultured in rich YE5S media until mid–log phase at 27°C and subsequently shifted to the restrictive temperature of 36°C before observation. To keep cultures at the proper temperature, a temperature-controlled chamber (Air Therm SMT; World Precision Instruments Inc., Sarasota, FL) was used. The sections of images acquired at each time point were compressed into a 2D projection using the DeltaVision maximum intensity algorithm. Deconvolution was applied before the 2D projection. Images were taken as 14–16 sections along the z axis at 0.2 µm intervals; they were then deconvolved and merged into a single projection. Captured images were processed with Photoshop CS6 (version 13.0; Adobe, San Jose, CA).

### Quantification of fluorescent signal intensities

For quantification of signal intensities of Klp9-GFP located at the spindle midzone during anaphase B, 14–16 sections were taken along the z-axis at 0.2-µm intervals. After deconvolution and projection images of maximum intensity, a 20 × 20–pixel (2.15-µm square) region of interest (ROI) with maximum sum intensity was determined. After subtracting the mean intensity of three regions around each ROI as background intensities, values of the maximum sum intensities were used for statistical data analysis.

## Statistical data analysis

We used the two-tailed unpaired Student’s *t*-test to evaluate the significance of differences in different strains. All the experiments were performed at least twice. Experiment sample numbers used for statistical testing were given in the corresponding figures and/or legends. We used this key for asterisk placeholders to indicate p-values in the figures: e.g., ****, P < 0.0001.

## Supporting information

## ACKNOWLEDGEMENTS

We are grateful to Paul Nurse, Iain Hagan, Jonathan Millar and J. Richard McIntosh for providing us with the strains used in this study. We thank Kylie Pan for critical reading of the manuscript and useful suggestions. This work was supported by the Japan Society for the Promotion of Science (JSPS) (KAKENHI Scientific Research (A) 16H02503 to T.T., a Challenging Exploratory Research grant 16K14672 to T.T., Scientific Research (C) 16K07694 to M.Y. and Scientific Research (C) 15KT0155 to K.F.), the Naito Foundation (T.T.) and the Uehara Memorial Foundation (T.T).

## AUTHOR CONTRIBUTIONS

M.Y. K.F. and T.T designed experiments. M.Y. K.F. M.O. and Y.T. performed experiments and analysed the data. T.T. organised a whole project. M.Y., K.F. and T.T. wrote the manuscript and M.O. and Y.T. made suggestions.

**COMPETEING INTERESTS:**
The authors declare no competing interests.

**ADDITIONAL INFORMATION:**
Supplementary information accompanies this paper at http://www.nature.com/srep.

## Abbreviations used

MAPs: microtubule associated proteins
NLS: nuclear localisation signal
SPB: spindle pole body
ts: temperature sensitive
TIRFM: total internal reflection fluorescence microscopy

