## Supplementary material for "Kinesin-6 Klp9 plays motor-dependent and -independent roles in collaboration with Kinesin-5 Cut7 and the microtubule crosslinker Ase1 in fission yeast"

**Supplementary Table S1:** Fission yeast strains used in this study

**Supplementary Figure S1:** Purification of the Klp9 protein for TIRF microscopy

**Supplementary Figure S2:** Genetic interactions between various *klp9* mutants and deletions of other MAP

### Supplementary Table S1: Fission yeast strains used in this study

| Strains | Genotypes | Figures used | Derivations |
|---|---|---|---|
| MO282 | $h^-$ *klp9-YFP-natR pcp1-CFP-hphR aur1R-Pnda3-mCherry-atb2 leu1 ura4* | 2A-B | This study |
| MO330 | $h^-$ $klp9^{rigor}$*-YFP-natR pcp1-CFP-hphR aur1R-Pnda3-mCherry-atb2 leu1 ura4* | 2A-B | This study |
| MO295 | $h^-$ *klp9-2-YFP-natR pcp1-CFP-hphR aur1R-Pnda3-mCherry-atb2 leu1 ura4* | 2A-B | This study |
| MO100 | $h^-$ *cut12-GFP-*$ura4^+$ *aur1R-Pnda3-mCherry-atb2 leu1 ura4* | 2C-E, 6D-F | This study |
| MO404 | $h^-$ *klp9::hphR cut12-GFP-*$ura4^+$ *aur1R-Pnda3-mCherry-atb2 leu1 ura4* | 2C-E, 6D-F | This study |
| MO389 | $h^-$ $klp9^{rigor}$*-kanR cut12-GFP-*$ura4^+$ *aur1R-Pnda3-mCherry-atb2 leu1 ura4* | 2C-E | This study |
| MO388 | $h^-$ *klp9-2-kanR cut12-GFP-*$ura4^+$ *aur1R-Pnda3-mCherry-atb2 leu1 ura4* | 2C-E | This study |
| MY984 | $h^-$ *klp9::hphR leu1 ura4* | 3A, 3C | This study |
| MO393 | $h^+$ *ase1::kanR leu1 ura4 his2* | 3A, 3C, 4A, 5A, 5D, S2B-C | This study |
| MO286 | $h^-$ $klp9^{rigor}$*-kanR leu1 ura4* | 3A-B | This study |
| MY1760 | $h^+$ *ase1::hphR leu1 ura4 his2* | 3A-B, 4A, S2B-C | This study |
| 513 | $h^-$ *leu1 ura4* | 3B | Our lab stock |
| MY1321 | $h^-$ *klp9-2-kanR leu1 ura4* | 3B | This study |
| MY1277 | $h^-$ $klp9^{rigor}$*-kanR ase1::hphR leu1 ura4* | 3B-C | This study |
| MY1220 | $h^-$ *klp9-2-kanR ase1::hphR leu1 ura4* | 3B-C | This study |
| MY990 | $h^+$ *cut7::bleR pkl1::natR leu1 ura4 his2* | 3B-C, S2A, S2C | This study |
| MO164 | $h^+$ *klp9-2-kanR cut7::bleR pkl1::natR leu1 ura4 his2* | 3B-C | This study |
| MY1462 | $h^-$ *klp9-GFP-kanR aur1R-Pnda3-mCherry-atb2 leu1 ura4* | 4A-C, 5A, 5C | This study |
| MY1735 | $h^-$ $klp9^{\Delta 38C}$*-GFP-kanR aur1R-Pnda3-mCherry-atb2 leu1 ura4* | 4A-C | This study |
| MO343 | $h^-$ $klp9^{\Delta 92C}$*-GFP-kanR aur1R-Pnda3-mCherry-atb2 leu1 ura4* | 4A-C | This study |
| MO365 | $h^-$ $klp9^{\Delta 133C}$*-GFP-kanR aur1R-Pnda3-mCherry-atb2 leu1 ura4* | 4A-C | This study |
| MO366 | $h^-$ $klp9^{\Delta 172C}$*-GFP-kanR aur1R-Pnda3-mCherry-atb2 leu1 ura4* | 4A, 4C | This study |
| MY1727 | $h^-$ $klp9^{\Delta 234C}$*-GFP-kanR aur1R-Pnda3-mCherry-atb2 leu1 ura4 lys1?* | 4A, 4C | This study |
| MO384 | $h^-$ $klp9^{KARAKA}$*-GFP-kanR aur1R-Pnda3-mCherry-atb2 leu1 ura4* | 4A-C | This study |
| MO362 | $h^-$ $klp9^{\Delta Motor}$*-GFP-kanR aur1R-Pnda3-mCherry-atb2 leu1 ura4* | 4A-C | This study |
| MY1331 | $h^-$ $klp9^{\Delta 38C}$*-GFP-hphR leu1 ura4* | 4A, S2B-C | This study |
| MO338 | $h^-$ $klp9^{\Delta 92C}$*-GFP-kanR leu1 ura4* | 4A, S2B-C | This study |
| MO319 | $h^-$ $klp9^{\Delta 133C}$*-GFP-kanR leu1 ura4* | 4A, S2C | This study |
| MO337 | $h^-$ $klp9^{\Delta 172C}$*-GFP-kanR leu1 ura4* | 4A, S2C | This study |
| MO316 | $h^-$ $klp9^{\Delta 234C}$*-GFP-kanR leu1 ura4* | 4A, S2C | This study |
| MO369 | $h^-$ $klp9^{KARAKA}$*-GFP-kanR leu1 ura4* | 4A, S2B-C | This study |
| MO368 | $h^-$ $klp9^{\Delta Motor}$*-GFP-kanR leu1 ura4* | 4A, S2B-C | This study |
| MO416 | $h^+$ $klp9^{\Delta CC1}$*-GFP-hphR aur1R-Pnda3-mCherry-atb2 leu1 ura4 his2* | 5A-C | This study |
| MO411 | $h^-$ $klp9^{\Delta CC2}$*-GFP-hphR aur1R-Pnda3-mCherry-atb2 leu1 ura4* | 5A-C | This study |
| MO405 | $h^-$ $klp9^{\Delta CC1}$*-GFP-hphR leu1 ura4* | 5A, 5D, S2C | This study |
| MO405 | $h^-$ $klp9^{\Delta CC2}$*-GFP-hphR leu1 ura4* | 5A, 5D, S2C | This study |
| YT317 | $h^-$ *cut7-GFP-kanR sid4-mRFP-natR aur1R-Pnda3-mCherry-atb2 leu1 ura4* | 6A, 6C | This study |
| YT080 | $h^+$ *cut7-122-GFP-kanR sid4-mRFP-natR aur1R-Pnda3-mCherry-atb2 leu1 ura4 his2* | 6A, 6D-G | This study |
| YT252 | $h^-$ *cut7-GFP-kanR klp9::hphR sid4-mRFP-natR aur1R-Pnda3-mCherry-atb2 leu1 ura4* | 6A | This study |

| | | | |
|---|---|---|---|
| YT074 | $h^{+}$ *cut7-122-GFP-kanR klp9::hphR sid4-mRFP-natR aur1R-Pnda3-mCherry-atb2 leu1 ura4 his2* | 6A, 6C-G | This study |
| YT344 | $h^{-}$ *cut7-GFP-kanR* $klp9^{rigor}$*-GFP-hphR aur1R-Pnda3-mCherry-atb2 leu1 ura4* | 6A | This study |
| YT352 | $h^{-}$ *cut7-122-GFP-kanR* $klp9^{rigor}$*-GFP-hphR aur1R-Pnda3-mCherry-atb2 leu1 ura4* | 6A, 6G | This study |
| YT287 | $h^{+}$ *cut7-GFP-kanR ase1::hphR sid4-mRFP-natR aur1R-Pnda3-mCherry-atb2 leu1 ura4* | 6A | This study |
| YT299 | $h^{+}$ *cut7-122-GFP-kanR ase1::hphR sid4-mRFP-natR aur1R-Pnda3-mCherry-atb2 leu1 ura4* | 6A, 6G | This study |
| MY1328 | $h^{-}$ $klp9^{\Delta 38C}$*-GFP-hphR pkl1::natR leu1 ura4* | S2A, S2C | This study |
| MO347 | $h^{-}$ $klp9^{\Delta 92C}$*-GFP-kanR pkl1::natR leu1 ura4* | S2A, S2C | This study |
| MO326 | $h^{-}$ $klp9^{\Delta 133C}$*-GFP-kanR pkl1::natR leu1 ura4* | S2C | This study |
| MO335 | $h^{-}$ $klp9^{\Delta 172C}$*-GFP-kanR pkl1::natR leu1 ura4* | S2C | This study |
| MO328 | $h^{-}$ $klp9^{\Delta 234C}$*-GFP-kanR pkl1::natR leu1 ura4* | S2C | This study |
| MY1762 | $h^{-}$ $klp9^{KARAKA}$*-GFP-kanR pkl1::natR leu1 ura4* | S2A, S2C | This study |
| MO359 | $h^{-}$ $klp9^{\Delta Motor}$*-GFP-kanR pkl1::natR leu1 ura4* | S2A, S2C | This study |
| MO417 | $h^{-}$ $klp9^{\Delta CC1}$*-GFP-hphR pkl1::natR leu1 ura4* | S2A, S2C | This study |
| MO421 | $h^{-}$ $klp9^{\Delta CC2}$*-GFP-hphR pkl1::natR leu1 ura4* | S2A, S2C | This study |

*Strains were developed for this study unless otherwise specified.
*his2=his2-245*; *leu1=leu1-32*; *ura4=ura4-D18*.

**Supplementary Figure S1. Yukawa *et al.***

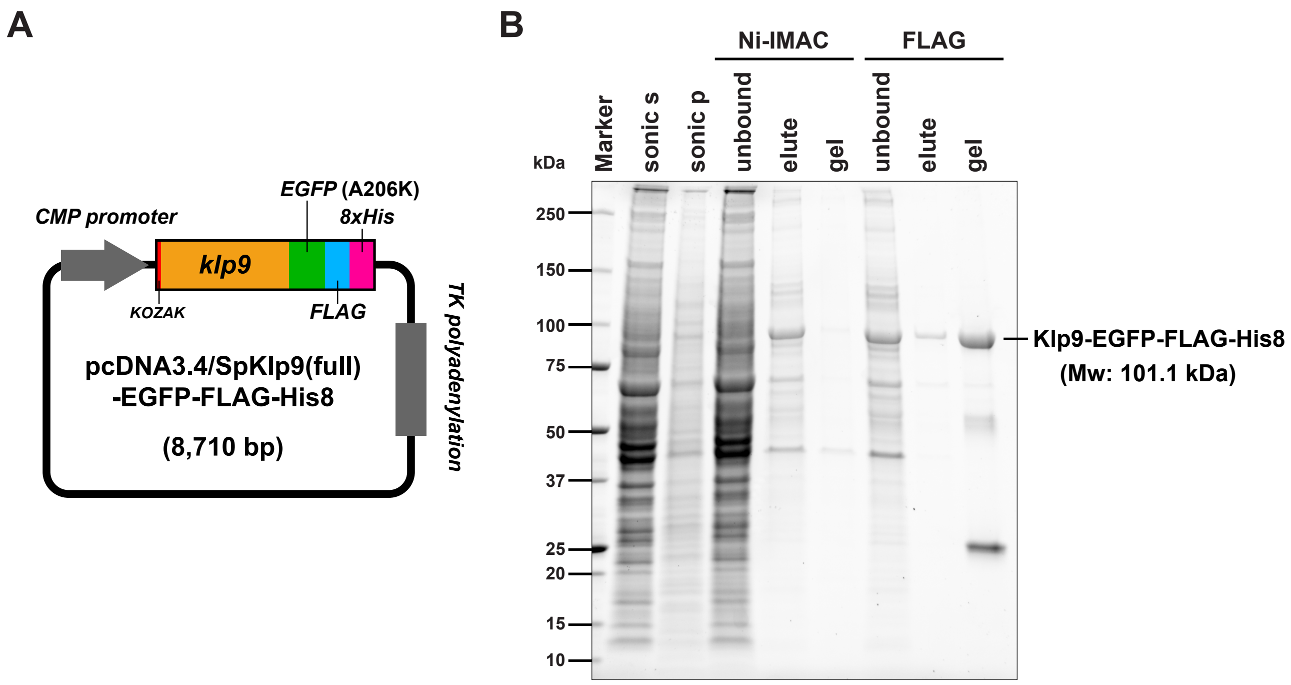


**Supplementary Figure S1. Purification of the Klp9 protein for TIRF microscopy**

**(A)** Diagram of a plasmid for transient expression full-length Klp9 in human cells.

**(B)** SDS-PAGE analysis (4-15%) of the full-length Klp9 protein. The protein bands were visualised by Stain-Free technology (Bio-rad). The right-most lane represents Klp9 preparations used for *in vitro* assays with TIRF.

**Figure S2. Yukawa *et al.***

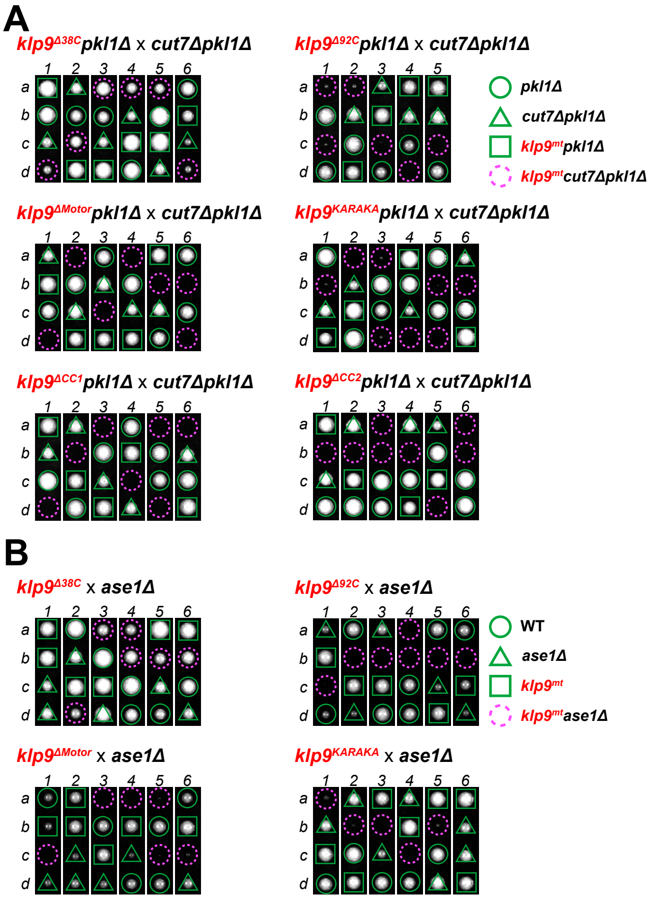


**C**

| | Synthetic lethality | |
|---|---|---|
| ***klp9* allele** | ***cut7Δpkl1Δ*** | ***ase1Δ*** |
| *klp9$^{+}$* | V | V |
| Δ38C | V | V |
| Δ92C | SL | SL |
| Δ133C | SL | SL |
| Δ172C | SL | SL |
| Δ234C | SL | SL |
| ΔMotor | SL | SL |
| KARAKA | SL | SL |
| ΔCC1 | SL | SL |
| ΔCC2 | SL | SL |

**Figure S2. Genetic interactions between various *klp9* mutants and deletions of other MAP**
**(A, B)** Tetrad analysis. Spores were dissected upon crosses between indicated strains Individual spores (a–d) in each ascus (1–8) were dissected on YE5S plates and incubated for 3 d at 27°C. Representative tetrad patterns are shown: images of dissected tetrads were merged, in which while spaces were created between each tetrad. **(A)** Circles, triangles and squares with green lines indicate *pkl1*Δ, *cut7*Δ*pkl1*Δ and *klp9$^{mt}$pkl1*Δ, respectively. **(B)** Circles, triangles and squares with green lines indicate wild type, *ase1*Δ, *klp9$^{mt}$*, respectively. Assuming 2:2 segregation of individual markers allows the identification of *klp9$^{mt}$cut7*Δ*pkl1*Δ **(A)** *klp9*Δ*ase1* Δ mutants **(B)** (indicated by dashed magenta circles). **(C)** A summary table indicating genetic interactions between various *klp9* mutants and *cut7*Δ*pkl1*Δ or *ase1*Δ. V stands for viable, while SL stands for synthetically lethal.
